# Development of the Inactivated QazCovid-in Vaccine: Protective Efficacy of the Vaccine in Syrian Hamsters

**DOI:** 10.1101/2021.07.13.452175

**Authors:** Kuandyk Zhugunissov, Kunsulu Zakarya, Berik Khairullin, Mukhit Orynbayev, Yergali Abduraimov, Markhabat Kassenov, Kulyaisan Sultankulova, Aslan Kerimbayev, Sergazy Nurabayev, Balzhan Myrzhakhmetova, Aziz Nakhanov, Ainur Nurpeisova, Olga Chervyakova, Nurika Assanzhanova, Yerbol Burashev, Muratbay Mambetaliev, Moldir Azanbekova, Syrym Kopeyev, Nurlan Kozhabergenov, Aisha Issabek, Moldir Tuyskanova, Lespek Kutumbetov

## Abstract

In March 2020, the first cases of human coronavirus infection COVID-19 were registered in Kazakhstan. We isolated the SARS-CoV-2 virus from the clinical material from the patients. Subsequently, a whole virion inactivated candidate vaccine, QazCovid-in, was developed based on this virus. To obtain the vaccine, a virus grown in Vero cell culture was used, which was inactivated with formaldehyde, purified, concentrated, sterilized by filtration, and then sorbed on aluminum hydroxide gel particles. The formula virus and adjuvant in buffer saline solution was used as a vaccine. The safety and protective effectiveness of the developed vaccine was studied on Syrian hamsters. The results of the studies showed the absolute safety of the candidate vaccine on the Syrian hamsters. When studying the protective effectiveness, the developed vaccine with an immunizing dose of 5 mcg/dose of a specific antigen protected animals from wild virus at a dose of 10^4.5^ TCID_50_/ml. The candidate vaccine formed virus-neutralizing antibodies in vaccinated hamsters in titers from 3.3 ± 1.45 log2 to 7.25 ± 0.78 log2, which were retained for 6 months (observation period) in the indicated titers. The candidate vaccine suppressed the replication of the wild virus in the body of vaccinated hamsters, protected against the development of acute pneumonia and ensured 100% survival of the animals. At the same time, no replicative virus was isolated from the lungs of vaccinated animals. At the same time, a virulent virus was isolated from the lungs of unvaccinated animals in relatively high titers, reaching 4.5 ± 0.7 lg TCID_50_/ml. After challenge infection, 100% of unvaccinated hamsters became ill with clinical signs (stress state, passivity, tousled coat, decreased body temperature and body weight, and the development of acute pneumonia), of which 25 ± 5% were fatal. The findings paved the way for testing the candidate vaccine in humans in clinical trials.

## Introduction

Coronaviruses are a large family of RNA-containing viruses that can infect humans and a number of animal species (Lau & Chan, 2015, p. 209; To et al, 2013, p. 103-108; Weiss & Navas-Martin, 2005). In humans, coronaviruses can cause a number of diseases-from a mild form of acute respiratory infection to severe acute respiratory syndrome (Weiss & Navas-Martin, 2005; Su et al., 2016). Currently, the following coronaviruses are known to circulate among the population: HCoV-229E, HCoV-OC43, HCoV-NL63, HCoV-HKU1, SARS-CoV, MERS-CoV, which, as a rule, cause diseases of the upper respiratory tract and lungs with moderate severity (Ye et al., 2020; Lim et al., 2016). A new coronavirus infection, COVID-19, caused by the SARS-CoV-2 virus, was added to this list, which was registered in the China at the end of December 2019 in Wuhan (Lu et al., 2020; Mackenzie & Smith, 2020). The new coronavirus infection has become a pandemic in a short time and has caused enormous socio-economic damage to the life and activities of humanity around the world. The rapid spread of the infection stopped many of the normal activities of entire states, and severe forms of the disease led to the death of many infected people as a result of the development of pneumonia (Lu et al., 2020). In order to prevent and combat the new coronavirus infection caused by the SARS-CoV-2 virus, many advanced countries of the world quickly began to create a means of specific prevention of the disease in the form of various types of vaccines. These types of vaccines being developed against the novel coronavirus infection COVID-19 include: RNA vaccines, DNA vaccines, recombinant vector vaccines, subunit vaccines, inactivated vaccines, and live vaccines (W.H.O., 2021; Krammer, 2020; Poland et al., 2020). The advantages and disadvantages of these vaccines are detailed in a number of literature sources (Dong et al. 2020; Li et al., 2020).

According to WHO data (as of May 7, 2021), more than 236 potential vaccines are being developed worldwide, of which 63 are being tested on humans (W.H.O., 2021). Of the latter, 26 candidate vaccines are undergoing phase III clinical trials, some of which are already being used for mass vaccination in a number of countries (W.H.O., 2021).

Since the beginning of the pandemic, Kazakhstan has also started developing a domestic vaccine against the new coronavirus infection on five platforms, of which one vaccine is based on the traditional technology of preparing an inactivated vaccine. This vaccine, under the name QazCovid-in, has successfully passed the I and II phase of clinical trials and is in the second half of the III phase of similar trials (Phase 1-2 results are under peer review in EClinicalMedicine journal).

In this article, we present the results of the creation of a new inactivated candidate vaccine QazCovid-in (or QazVac) and the study of its safety and immunological effectiveness in Syrian hamsters.

## Materials and methods

### Virus

In the research on the development of the QazCovid-in vaccine against COVID-19 coronavirus infection, the SARS-CoV-2/KZ_Almaty04.2020 strain (Kutumbetov et al., 2020) deposited in the republican depository of the collection of microorganisms of the RSE “Research Institute for Biological Safety Problems” (RIBSP) of the Committee of Science of the Ministry of Education and Science of the Republic of Kazakhstan, obtained from the SARS-CoV-2 virus isolated from a clinical sample of a patient with COVID-19 coronavirus infection, was used. The virus strain was partially sequenced and the nucleotide sequences of its E, N, S, and ORF1a genes were determined. The results of virus isolation from clinical material and sequencing of these genes are presented in supplementary material 1.

### Animals

The experiments used 100 of Syrian hamsters, randomly selected by randomization. The hamsters were kept in individual ventilated complexes of the Delta IVC-ZJ3 brand (China).^1^ The experimental animals were monitored daily with the cataloguing of their general clinical condition, body temperature, body weight, appetite and water intake.

### Preparation of the candidate vaccine

The virus was cultured in Vero cells (WHO, Lot No. CB0 or CB884) at a temperature of 37 °C for 48 hours, and then inactivated with formaldehyde (Sigma Aldrich) for 24 hours. The completeness of inactivation of the virus was established by a bioassay in a cell culture carried out in three passages. The inactivated virus was purified and concentrated by a combined method using tangential flow ultra-filtration and exclusive chromatography. The purified and concentrated suspension of the inactivated virus was sterilized by filtration through membrane filters with a pore diameter of 0.22 microns. The purified virus, taken at a certain concentration of a specific protein in a phosphate-salt buffer, was mixed with an Al (OH)_3_ adjuvant (Alhydrogel®, InvivoGen, France) and used as a candidate vaccine against SARS-CoV-2. The technology of preparation of the candidate vaccine is schematically illustrated in Fig. 1.

**Figure 1.**
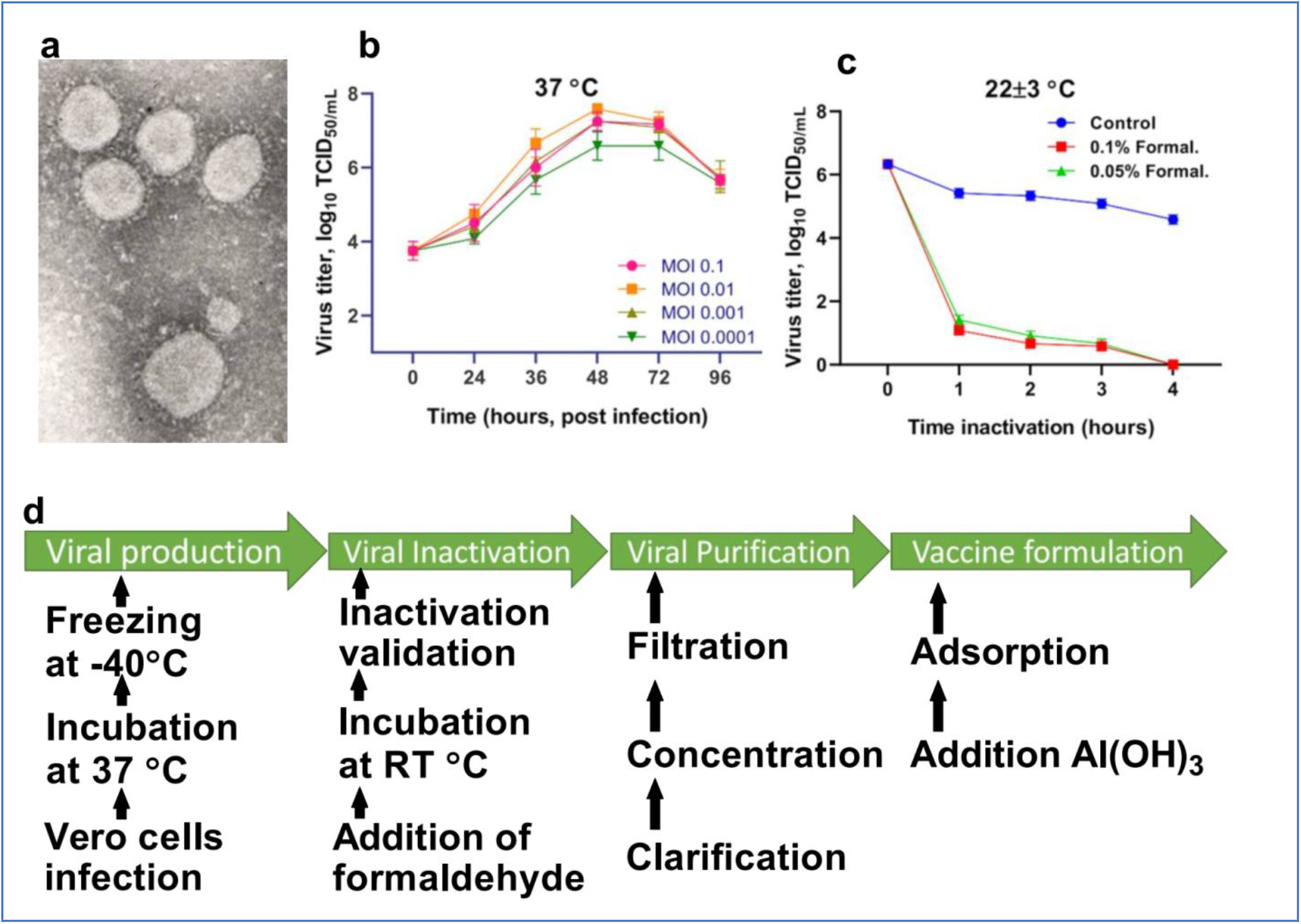
Scheme of preparation of the candidate vaccine QazCovid-in. Visual **(a)** shows an electronic micrograph of the SARS-CoV-2 virus (increased by x18000). During the study of the SARS-CoV-2 virus by electron microscopy, the morphological features of the coronavirus virions were determined, which are represented by a spherical shape with dimensions from 80 to 200 nanometers and containing club-shaped glycoprotein peplomers with dimensions of 15-20 nanometers. In the studied population of coronavirus, virions with a size of 120 - 125 nanometers predominate most of all. The virus was isolated in a Vero cell culture from clinical samples taken from patients of the Republic of Kazakhstan (Almaty) who were infected with the new COVID-19 coronavirus infection. The virus was identified by molecular genetic methods and identified as the SARS-CoV-2 coronavirus. Visual **(b)** shows the accumulation of the virus in Vero cells, depending on the multiplicity of the infecting dose (MOI). At the same time, the highest titer of the virus was detected in the period from 30 h to 42 h at a MOI of 0.1-0.01 TCID_50_/cell. However, no significant difference was found between the virus titers at different MOI (p≥0.05). Visual **(c)** shows the inactivation of the virus at room temperature. When determining the completeness of inactivation, the virus treated with formaldehyde did not cause cytopathic effect (CPE) in the monolayer of Vero cell culture with three passages every 4-5 days. Visual **(d)** shows the preparation steps for the QazCovid-in vaccine.

### Vaccine safety assessment

10 Syrian hamsters were administered the vaccine intramuscularly once in a dose of 0.5 ml, containing 5 μg of a specific virus protein and 1.0 mg of aluminum hydroxide on a phosphate-buffer saline solution (PBS). The control group of animals, which also consisted of 10 heads, was injected with a PBS intramuscularly in a volume of 0.5 ml. The animals were monitored daily for 20 days, with body temperature measured and live body weight recorded. The day before vaccination and 21 days after the introduction of the vaccine, blood samples were taken from animals of both groups, which were subjected to biochemical and hematological analysis. The scheme of the safety of the vaccine is shown in Fig. 2a.

**Figure 2.**
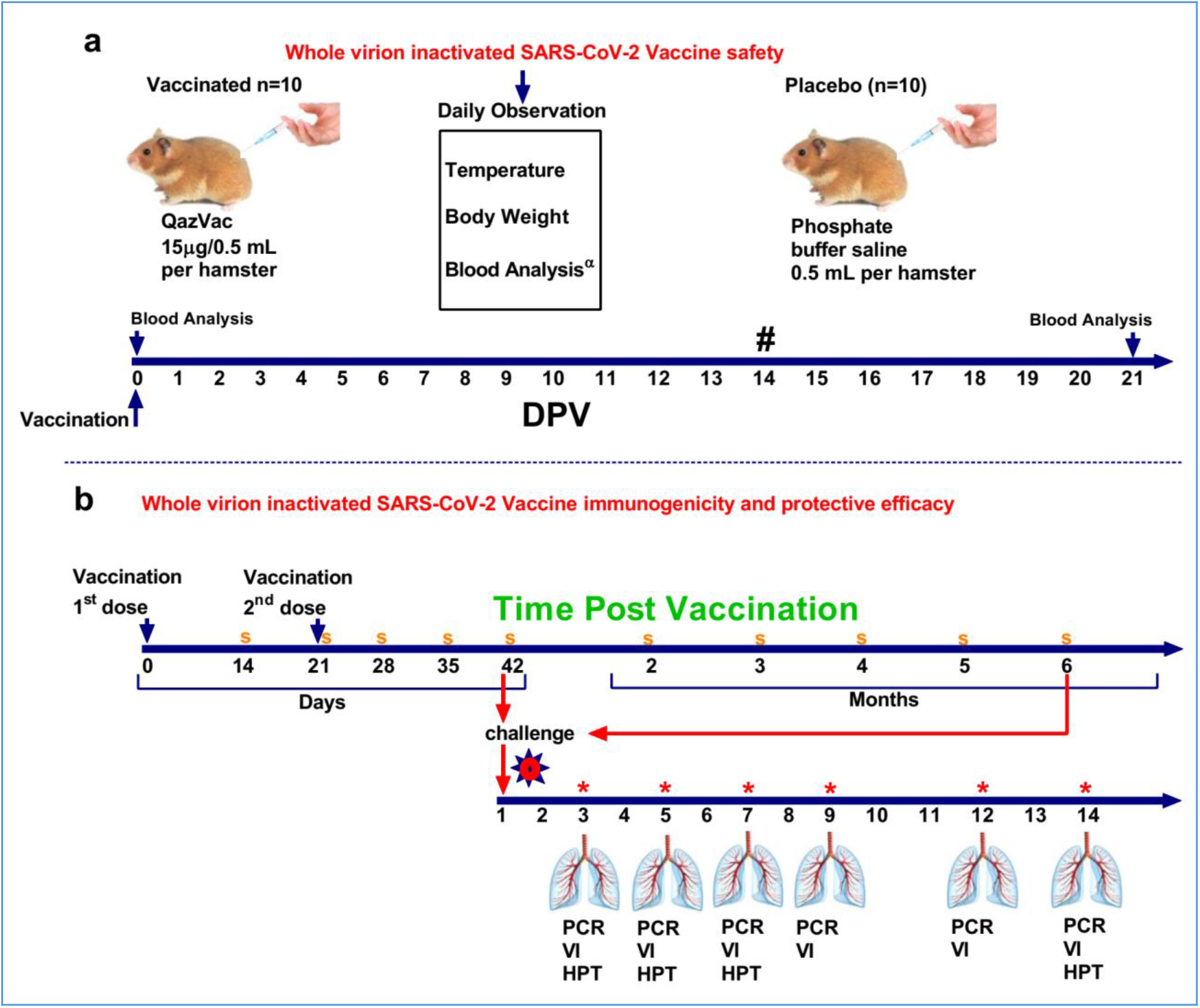
Design of the study on Syrian hamsters. Part (**a)** shows the study of the safety of the inactivated QazCovid-in vaccine in Syrian hamsters. The safety study was conducted on 2 groups of animals. The first group of animals (n=10) was vaccinated with the inactivated QazCovid-in vaccine at a dose of 15μg/0.5 ml/head intramuscularly. The second group of animals (n=10) received a PBS in the volume of 0.5 ml/head also intramuscularly. The animals were monitored on a daily basis, with body temperature and body weight measured. Hematological and biochemical blood tests were performed before and on the 21st day post vaccination (DPV). Measurement of body temperature and body weight was carried out within 14 days. Part (**b)** shows the determination of immunogenicity and evaluation of the protective efficacy of the inactivated QazCovid-in vaccine in Syrian hamsters. The first dose of the vaccine was administered on day 0, the second dose, on DPV 21. Sampling (**s**) of animal blood serum on 14, 21, 28, 35, 42 DPV and 2, 3, 4, 5, 6-months after vaccination to determine the VNA in the neutralization test. On the 42^nd^ and 180^th^ (6 months) DPV, challenge infection with the wild SARS-CoV-2 virus was carried out at a dose of 10^4.5^ TCID_50_/head in the volume of 100 μl intranasally, 50 μl in each nostril. On the 3rd (n=5), 5th (n=5), 7th (n=5), 9th (n=5), 12th (n=5), 14th (n=5) day post infection (DPI), lung samples were taken by biopsy with determination of viral load by rtPCR, viral release on Vero cell culture and histopathology. **PCR** - polymerase chain reaction.**VI** - virus isolation. **HPT**- histopathology. Note (*) - on 3, 5, 7, 9, 12, 14 DPI samples of nasal and oral swabs were collected and examined by PCR and viral isolation in cell culture.

### Hematological and biochemical blood tests

Hematological analysis of blood samples was performed using an automatic blood analyzer T-540 Coulter (Coulter Electronics, Hialeah, FL, USA). As a result of the analysis, the concentration of hemoglobin, hematocrit, red blood cells, white blood cells, platelets, neutrophils, eosinophils, lymphocytes and monocytes was determined. Biochemical studies of blood serum samples were carried out on VITALAB SELECTRA 2 in an automatic analyzer (Merck, Germany) using commercial kits (DIASYS Diagnostic Systems GmbH, Germany), which determined the quantitative parameters of total protein, total bilirubin, glucose, creatinine, aspartate aminotransferase and alanine aminotransferase.

### Vaccine immunogenicity analyses and Protection study

60 Hamsters were administered the vaccine twice, intramuscularly, with an interval of 21 days, in a dose of 0.5 ml containing 5 μg of a specific virus protein adsorbed on aluminum hydroxide. The control group of animals in the amount of 20 heads of an analogous type was injected with a PBS in the volume of 0.5 ml, also intramuscularly. The animals were monitored on a daily basis, with body temperature measurement and live weight determination using electronic scales for 42 days. The day before the introduction of the vaccine, then on the 14th, 21st, 28th, 35th, 42nd day and every subsequent month after the introduction of the first dose of the vaccine for 6 months, blood serum samples were collected from all experimental animals to establish the dynamics of the formation of virus neutralizing antibodies (VNA) in the neutralization test.

To establish the protectivity of the vaccine, the infectious process was modeled on vaccinated and control (unvaccinated) Syrian hamsters by injecting them with the wild SARS-CoV-2 virus at a dose of 10^4.5^ TCID_50_/head in the volume of 100 μl intranasally. To do this, 50 % of vaccinated and unvaccinated Syrian hamsters were exposed to a virulent virus 42 days after the first vaccination, and the remaining 50% of vaccinated and control animals were exposed to a virulent virus after 6 months.

After infection with a virulent virus, the animals were monitored daily, recording their general condition, appetite, body temperature, live weight, and signs of pathologies. On days 3, 5, 7, 9, 12, and 14, samples of oral and nasal swabs were collected to determine the presence of a virulent pathogen.

To evaluate the effectiveness of the candidate vaccine in intramuscular administration of animals, 3 (n=5), 5 (n=5), 7 (n=5), 9 (n=5), 12 (n=5) and 14 (n=5) days after the challenge infection, the animals were euthanized by Carbon dioxide inhalation using a standard two-cell kit AE0904. The gas consumption was 3.5 l / min per cell for 2-4 minutes. The onset of animal death was monitored by the absence of respiration and fading of the eyes of each animal, after which a visual examination, autopsy, and lungs were selected for virus isolation and histological examination. The design of the study is shown in Fig. 2b.

### Histological examination of the lungs

For microscopic analysis, lung tissue samples were taken from all the studied animals and recorded in a 10% solution of neutral formalin. The tissue pieces were left in formalin overnight at room temperature, then treated according to the standard histological technique procedure (dehydration, clearing, and compaction). Tissue sections with a thickness of 4-5 microns were prepared from paraffin blocks using a sled microtome. For a general overview, the histological sections are stained with hematoxylin and eosin. Microscopic analysis and photography was carried out under a Nikon ECLIPSE 50i microscope equipped with a Nikon Digital Sight DS-Fi1 camera (Japan).

### Neutralizing assay

Serum samples collected from immunized animals were inactivated at 56°C for 0.5h and serially diluted with cell culture medium in two-fold steps. The diluted serums were mixed with a virus suspension of 100 TCID_50_ in 96-well plates at a ratio of 1:1, followed by 2 hours incubation at 37 °C in a 5% CO2 incubator. 1-2×10^4^ Vero cells were then added to the serum-virus mixture, and the plates were incubated for 5 days at 37 °C in a 5% CO_2_ incubator. Cytopathic effect (CPE) of each well was recorded under microscopes, and the neutralizing titer was calculated by the dilution number of 50% protective condition.

### Virus RNA isolation

The virus RNA was extracted from clinical samples using the QIAamp viral RNA mini kit (QIAGEN, Hilden, Germany) according to the manufacturer’s instructions.

### Viral RNA analysis

SARS-CoV-2 RNA was assessed by RT-PCR using an approach similar to previously described (Corman et al., 2020; Wölfel et al., 2020). The following primers and probe were used to amplify the N gene of the SARS-CoV-2 virus: N_Sarbeco_F(cacattggcacccgcaatc), N_Sarbeco_R (gaggaacgagaagaggcttg), and N_Sarbeco_P (fam-acttcctcaaggaacaacattgcca-bbq) (Corman et al., 2020). The viral genome was evaluated by quantitative real-time PCR using the Superscript ® III Platinum ONE-STEP RT-PCR system kit with the Platinum™ Taq DNA polymerase system (Invitrogen, USA) according to the manufacturer’s instructions. The reactions were carried out in a thermal cycler of the Rotor-Gene 6000 series (Qiagen, Germany) with the following program: 1 reverse transcription cycle at 50°C for 20 min., 1 cycle of 95°C for 3 min followed by 45 cycles of 95°C for 15 s, 58°C for 30 s.

### Isolation of the virus in cell culture

The virus was isolated from lung samples in which viral RNA was detected by PCR. For this purpose, a 20% organ-tissue suspension was prepared from the lungs of hamsters using a generally accepted technique. Before infection, all mattresses with cell culture were microscopically examined and only mattresses with a good, typical monolayer were selected. After removing the culture medium, the prepared 20% suspension in the volume of 0.5 ml was applied to the monolayer of Vero cell culture and kept for 60 minutes at a temperature of 37 °C. Then the inoculate was removed, the monolayer was washed in three shifts with PBS solution, DMEM maintenance medium was added with fetal blood serum, and cultivation was continued at 37 °C with daily microscopy of the cell culture monolayer. The presence of the virus was determined by the cytopathic effect in infected cell cultures compared to the control uninfected cell culture. In the absence of cytopathic effect in a Vero cell culture infected with biomaterial samples, “blind” passaging was performed for at least three generations.

### Facility and ethics statements

Animal challenged experiments and all other experiments with live SARS-CoV-2 virus were housed under ABSL-3 conditions and BSL-3 facilities in the RIBSP. This study was performed in compliance with national and international laws and guidelines on animal handling, and the experimental protocol was approved by the Committee on the Ethics of Animal Experiments of the RIBSP of the Science Committee of the Ministry of Education and Science of the Republic of Kazakhstan (permit number: KZ0520/013, KZ1120/014).

### Statistics

Statistical analysis was performed using Graph Pad Prism 8.4.2 (GraphPad Software) to evaluate *p-values* for the overall experiment for vaccinated and control groups followed by the Student parametric test and the Wilcoxon-Mann-Whitney non-parametric test. One of the criteria for evaluating the effectiveness is the percentage of survival (PV). PV was evaluated by the Kaplan-Meyer method, and PV indicators in the groups of vaccinated and unvaccinated animals were compared by the log-rank method. P-values of less than .05 were considered significant.

## Results

### Safety of the QazCovid-in vaccine candidate on hamsters

During the entire observation period, the general condition of the animals remained satisfactory. They moved freely and actively in the cages, ate food well and took water. No pathologies of a general or local nature were observed in the animals. No changes in the growth and development of the animals were detected. The temperature response remained within the physiological norm. The obtained data of clinical observation indicate that the candidate vaccine, when administered intramuscularly at a dose of 15μg/0,5mL/head, did not have a negative effect on the overall clinical condition (behavior, appetite, etc.) of the test animals during the entire observation period. Similar data were obtained in studies in which Syrian hamsters were injected with a 3-fold dose of the test vaccine to detect allergenic and toxic properties. None of the 10 animals vaccinated with an excessive dose of the vaccine showed any signs of the disease during the entire period of clinical observation. The data obtained showed the absence of local and local irritant effects of the vaccine, as well as the ability of the drug to induce allergic reactions of immediate and delayed types, which minimizes the risks of anaphylactic reactions (anaphylactic shock, edema, etc.) to the administration of the vaccine.

### Hematological and biochemical studies

Hematological and biochemical blood tests showed that all the studied parameters of the animals ‘blood remained within the physiological norm during the entire follow-up period (Tables 1, 2). The hematological and biochemical parameters of the animals’ blood after immunization did not significantly differ from their background values in all groups (p≥ 0.05) established before vaccination. In experimental animals, slight fluctuations in the analyzed blood parameters were observed, but no definite trend in blood parameters was detected during the observation process.

**Table 1-.**
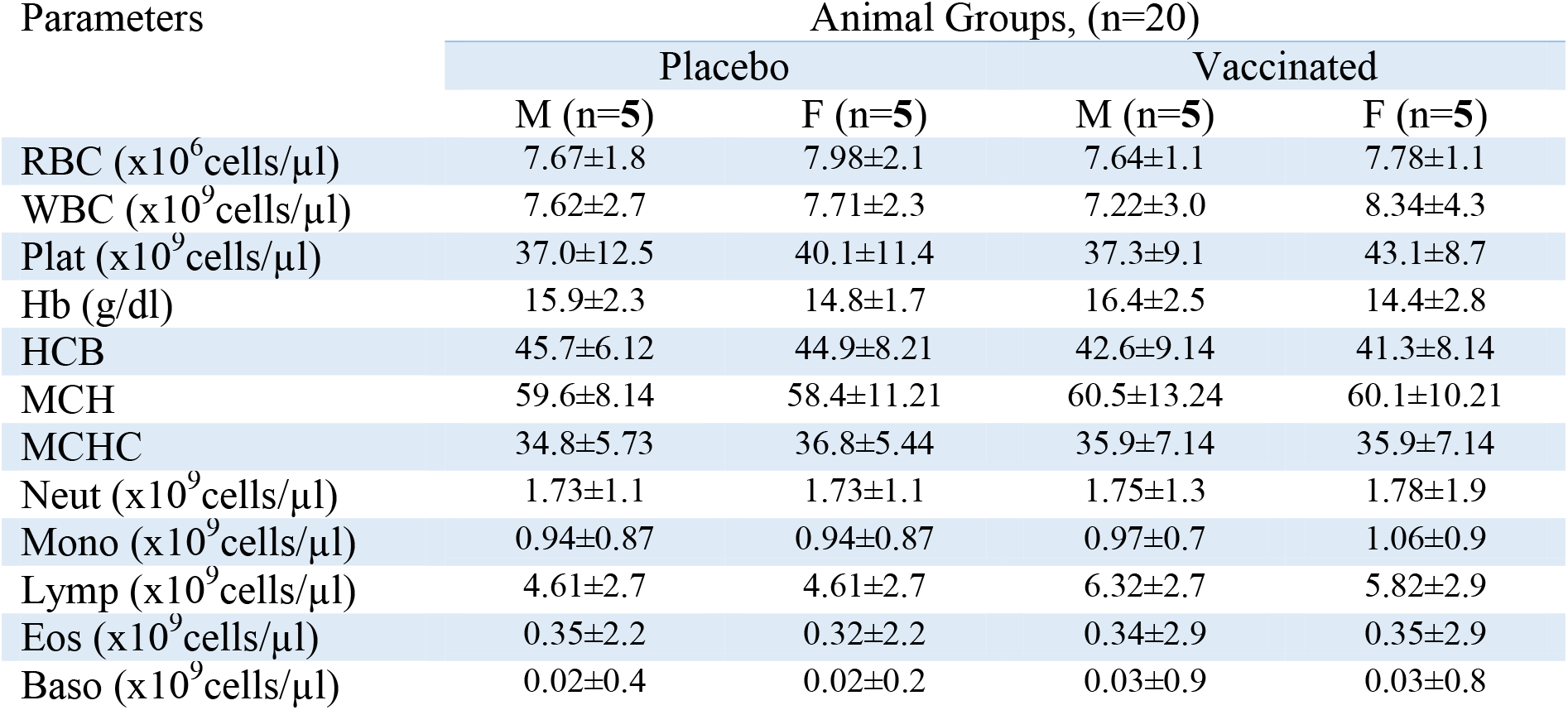
Average values (M±SD), hematological blood analysis of vaccinated hamsters after administration of the QazCovid-in vaccine candidate.

**Table 2-.** Average values (M±SD) of the main biochemical parameters of hamster blood serum in determining the safety of the test drug.

| Parameters | Animal Groups, (n=20) |  |  |  |
| --- | --- | --- | --- | --- |
|  | Placebo |  | Vaccinated |  |
|  | M (n=5) | F (n=5) | M (n=5) | F (n=5) |
| Total Protein (mg/dl) | 79.5±8.8 | 79.4±6.9 | 78.3±7.8 | 81.4±7.6 |
| Glucose (mg/dl) | 8.0±1.4 | 7.9±0.3 | 8.0±0.9 | 8.0±1.8 |
| Creatinine(mg/dL) | 66.9±3.8 | 66.7±3.9 | 67.0±3.2 | 67.3±3.7 |
| Total Bilirubin(mg/dl) | 9,2±1,1 | 8.6±1.7 | 7.8±0.6 | 7.9±0.7 |
| AST (U/L) | 97.8±25.2 | 93.5±14.8 | 97.9±22.3 | 94.1±23.0 |
| ALT (U/L) | 71.4±8.9 | 72.1±11.2 | 72.5±8.2 | 71.1±11.2 |
| Urea (mg/dL) | 4.9±0.9 | 4.8±0.8 | 4.8±0.2 | 4.9±0.3 |

### Antibody Responses after QazCovid-in Vaccination

The QazCovid-in vaccine in the body of vaccinated hamsters stimulated the formation of humoral immunity factors with a pronounced level (Fig. 3a). After the first dose of the vaccine was administered on day 14, VNA was detected in animal serum samples at a titer of 1.0±0.5 (CI 95%, <0.5-2.0) log_2_. On day 21, the titers of these antibodies increased to 4.8±0.9 (CI 95%, <3.0-6.0) log_2_. On the 28 days after the first dose of the vaccine, the level of antibodies increased slightly (CI 95%, 5.5±0.5 log_2_ (CI 95%, <5.0-7.0)), and on the 35 days after the first dose, their titer reached the highest values (7.25±0.78 log_2_ (CI 95%, <7.0-8.0)), and remained at this level until the 42nd day. A decrease in the VNA titer was observed at 2 months after the second dose (6.7±0.7 log_2_ (CI 95%, <6.0-8.0)), and at 6 months, the antibody titer averaged 3.3±1.45 (CI 95%, <2.0-7.0) log_2_. The analysis of the obtained research results shows that the level of VNA in the blood serum samples of immunized hamsters between the first and second vaccinations had a significant difference (P≤ 0.001).

**Figure 3.**
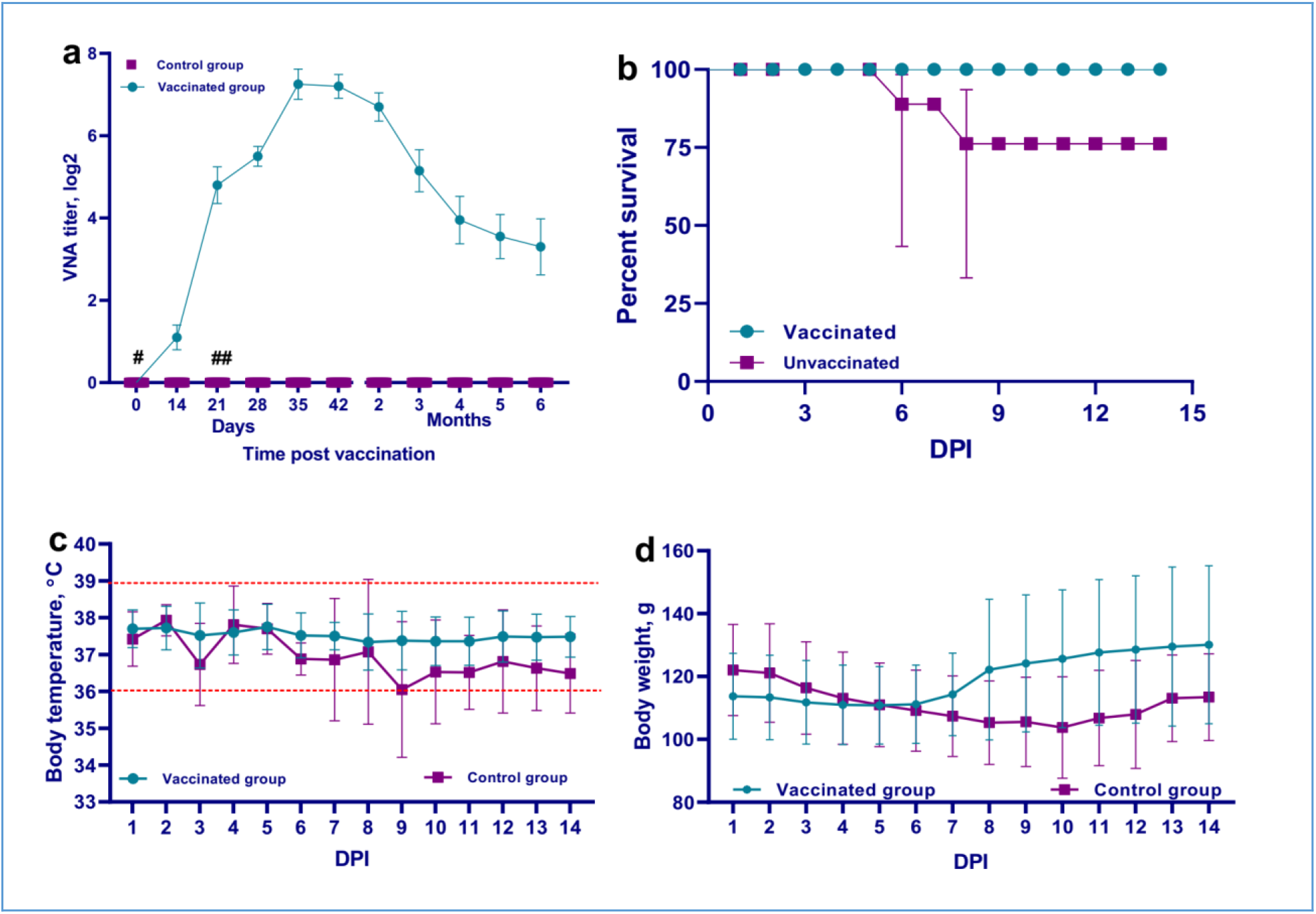
Performance indicators of the inactivated QazCovid-in. Part (**a)** shows vaccine-VNA in blood serum samples of Syrian hamsters before vaccination (day 0) and after the first (days 14, 21), second vaccination (days 28, 42 and months 2, 3, 4, 5, 6). The second vaccination was carried out on day 21. The graph shows the average values of VNA titers with standard deviation (M±SD). (* * *) – p < 0.001. A statistically- significant increase in VNA titers was shown for the vaccinated group on days 14 and 21 after the first immunization compared to days 7, 14 and 21 after the second immunization. The survival rate (**b**) of vaccinated and unvaccinated hamsters, after infection with the wild SARS-CoV-2 virus, shows the effect of vaccination with the inactivated QazCovid-in vaccine. In the group of vaccinated animals, there were no cases of disease and death after infection with the SARS-CoV-2 virus, while in unvaccinated animals, stress, shortness of breath, passivity, a decrease in body temperature to 34.2°C, live weight and death were noted. In the lung tissue of the fallen animals, diffuse compaction, atelectasis, fullness, edema, and focal hemorrhage were noted. Mortality in the control group averaged 25±5% with 100% morbidity. The survival rate of animals after control infection was 100% in the vaccinated group and 75 % in the unvaccinated group (±95% CI). However, the differences in survival between the groups were statistically insignificant, χ^2^=1.2, p= 0.277. The temperature reaction (**c**) of vaccinated and unvaccinated hamsters after challenge with wild SARS-CoV-2 virus was noted. According to some literature sources, the limit of the normal body temperature of hamsters is from 36.2 °C to 37.5 ° C (Dutta & Sengupta., 2019). Based on the literature data and our own data obtained during the control of the body temperature of healthy hamsters, the body temperature of the used laboratory animals was taken to be from 36.0 ° C to 38.9°C, outside the physiological norm of the body temperature. In our studies, body temperature above 39.0 °C and below 35.5 °C was observed only among sick hamsters. The body temperature of vaccinated hamsters ranged from 37.0 °C to 38.5 °C, and non-vaccinated hamsters ranged from 34.2 °C to 39.0°C. The average body temperature values of the vaccinated and unvaccinated groups of animals did not have significant differences (p?0.05). Changes in the body weight (**d**) of vaccinated and unvaccinated hamsters after challenge with the wild SARS-CoV-2 virus were noted. There was a progressive decrease in the body weight of unvaccinated animals throughout the entire observation period. The disease of sedi infected control animals ended in a 25% mortality rate. While among animals immunized with the inactivated QazCovid-in vaccine, there were no signs of the disease or of a decrease in live weight or of mortality after infection with the virulent virus.

### Protective Efficacy of QazCovid-in Vaccine Candidates

After the challenge infection, the vaccinated group of hamsters completely lacked any signs of the disease, and all vaccinated animals remained alive and clinically healthy for 14 days. Their body temperature fluctuated within the physiological norm (36.0-39.0 °C) (Fig. 3c). However, in the first 5-6 days after infection with the virulent virus, there was no gain in the live weight of vaccinated hamsters, and from day 7 it began to grow actively until the end of the observation. The live weight of the vaccinated hamsters at the end of the observation did not differ from the live weight of similar animals of the control group that was not vaccinated and not infected (Fig. 3d). At the same time, in the animals of the unvaccinated group, there was a stress state, passivity, ruffled hair and stroking of the nasal mirror with the limbs, which was a sign of itching. Starting on the 3rd DPI, some unvaccinated animals showed a decrease in body temperature to 34.5°C, as well as a decrease in live weight, but there were no significant differences between the groups (p≥0.05). The peak of the disease occurred on 5-8 DPI. The death of unvaccinated animals occurred on the 3rd DPI. At the same time, 2 hamster died on the 3rd DPI, 2 more hamster on the 4th day, and 1 head fell on the 6th and 8th days. In unvaccinated animals, stress, passivity, moisture and tousled coat and rubbing of the noses by limbs were noted. Mortality ranged from 25 ± 5% to 35 ± 15% (± 95% CI) (Fig. 3b).

### Isolation of virus RNA from nasal and oral swabs from vaccinated and unvaccinated animals after control infection

In all samples (oral and nasal swabs) obtained from unvaccinated animals, SARS-CoV-2 virus RNA was detected (Ct<12.3-29.7) at all studied time points after the challenge infection. In vaccinated animals, RNA of the same virus was also detected in nasal swabs (Ct<22.3-26.8) collected from day 3 to day 9 and in oral swabs (Ct<23.0-27.5) collected from day 3 to day 5 (Fig. 4a and 4c). The remaining samples collected from the oral and nasal cavities up to the 14th day showed negative results (Ct<33.3-37.5). To detect the presence of a reproductive (live) infectious virus in the respiratory organs of hamsters, the virus was simultaneously isolated in cell culture from samples of nasal and oral swabs.

**Figure 4.**
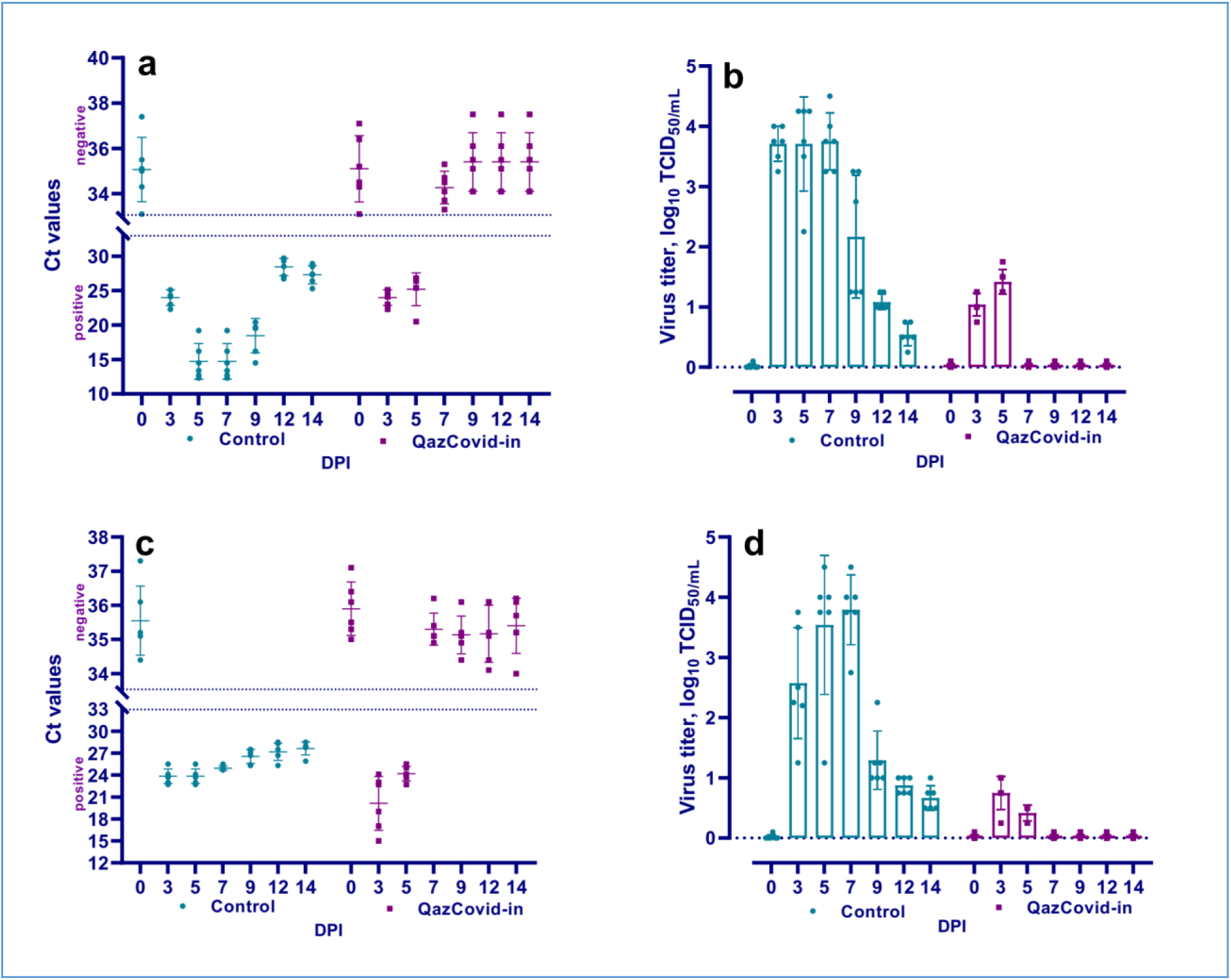
**describes** the nasal swabs PCR results (a), the oral swabs PCR data (c), and the virus isolation in cell culture from clinical samples from vaccinated and unvaccinated hamsters after control infection with wild SARS-CoV-2 virus (b –nasal swabs; d – oral swabs). Ct values between 10 and 33 are positive, while Ct values ≥33.1 are negative.

### Isolation of the virus in cell culture from nasal and oral swabs from vaccinated and unvaccinated animals after challenge

The virus was isolated from all nasal and oral samples collected from unvaccinated animals from 3 to 14 DPI, with a titer of 0.54±0.10 to 3.79±0.23 lg TCID_50_/ml. In studies of vaccinated animals, this pathogen was isolated from samples collected from 3 to 5 DPI, with a significantly low titer compared to the titer of the virus isolated from unvaccinated animals (Fig. 4b. 4d). At the same time, the difference between the titers of the virus isolated in the two groups of animals had significant differences (from p≤ .002 to p≤ .0001).

### Pathology and Viral Load in Lung Tissue after Challenge

There were no visible pathologies in the abdominal organs of the killed and fallen hamsters on the 3rd, 5th, 7th, 9th, 12th and 14th DPI of the vaccinated and unvaccinated hamsters, but when opening the chest cavity in unvaccinated hamsters, uneven staining and spot hemorrhages in the lungs were recorded. The bronchial and mediastinal lymph nodes were enlarged and swollen. In the other internal organs, no visible pathologies were detected. In vaccinated animals, the organs of the thoracic cavity, as well as the abdominal cavity, had no visible changes.

Further, the presence or absence of the viral genome in the upper and lower respiratory tracts of vaccinated and unvaccinated animals in the above-mentioned periods after control infection was established by PCR (Fig. 5). As a result, it was revealed that the genomes of the SARS-CoV-2 virus were present in the upper and lower respiratory tracts of both vaccinated and unvaccinated hamsters at the studied time (Fig. 5a). At the same time, the concentration of the virus genome in the vaccinated group of hamsters (Ct 24.20-36.80) was significantly lower than in the unvaccinated group of animals (Ct 11.53-30.10).

**Figure 5.**
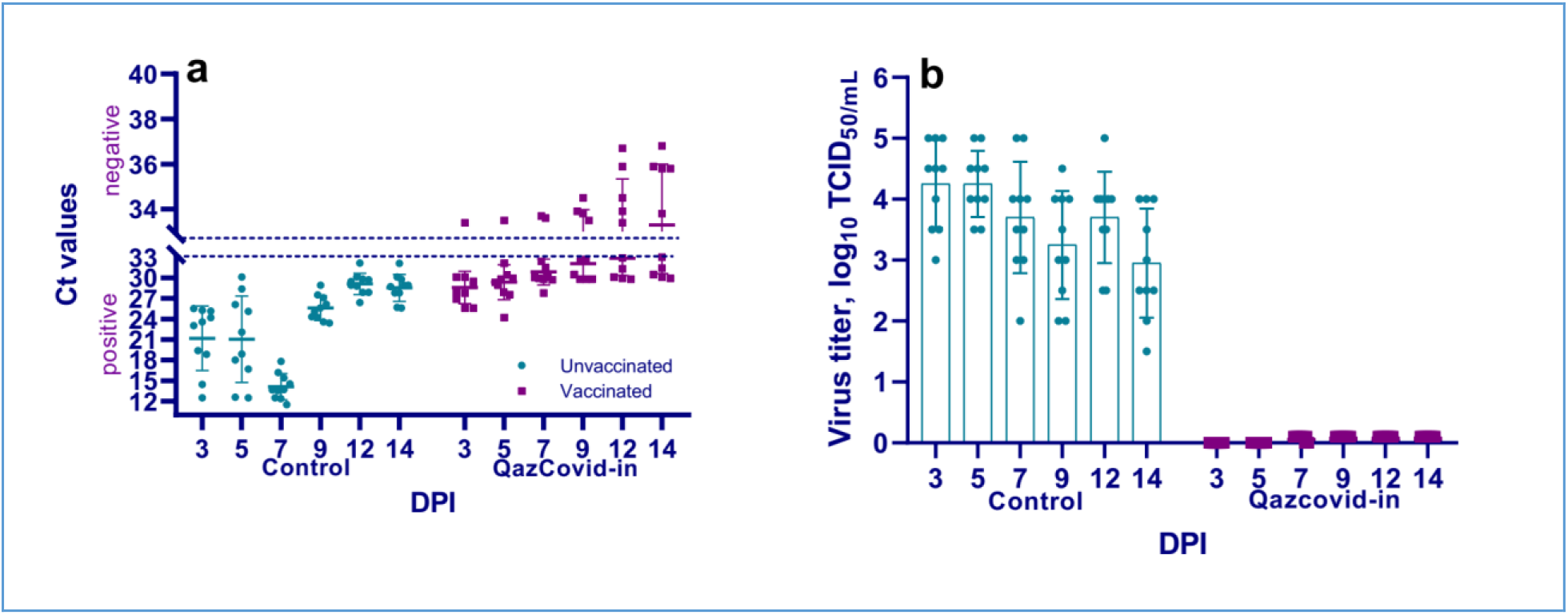
The presence and concentration of virulent virus in the lungs of vaccinated and unvaccinated hamsters. This figure shows data from the 3rd, 5th, 7th, 9th, 12th and 14th DPI with the wild SARS-CoV-2 virus, specifically the genomic RNA in the lungs (a), and virus isolation from the lungs in Vero cells (b). Ct values between 10 and 33 are positive, while Ct values ≥33.1 are negative.

In the case of a cell culture conducted to detect the presence of a replicative virus in respiratory tract tissue samples obtained from vaccinated animals, the results of virus isolation were negative, indicating the absence of an infectious SARS-CoV-2 virus capable of replication. In contrast to these data, in unvaccinated animals the virulent virus in the lung tissues was detected in relatively high titers, reaching 4.5±0.7 lg TCID_50_/ml (Fig.5c).

### Lung tissue histopathology of vaccinated and unvaccinated Syrian hamster after challenge

No visible morpho-pathological changes were detected in any internal organs, except for the lungs, of the animals used in the experiments.

Histological analysis after inoculation with a virulent virus in the lungs of animals of the unvaccinated group revealed chronic interstitial inflammatory cells during all the study periods (Fig. 6a, 6b, 6c, 6d). Histopathology in the lungs of animals of the vaccinated group was limited to minor alveolar changes which was detected only on the 3rd and 5th DPI with the virus, and they were less noticeable compared to the group of unvaccinated animals (Fig. 6e, 6f,). On the 7th, 9th, and 14th DPI, no changes in the lungs were observed in the animals of the vaccinated group (Fig. 6j, 6h).

**Figure 6.**
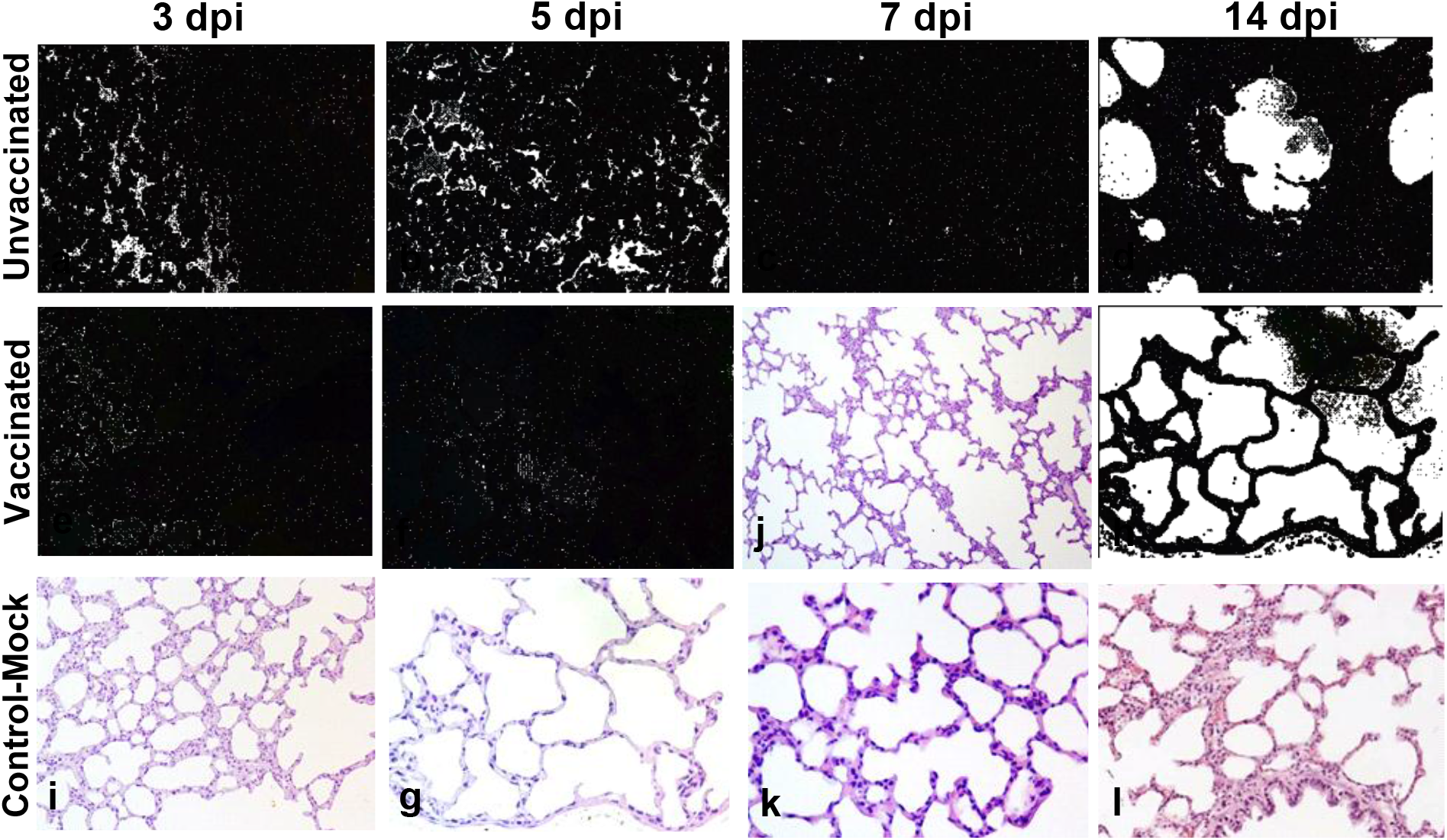
Paraffin sections of the lungs of vaccinated and unvaccinated hamsters. Figure 6 shows a pathohistological picture of the lung of unvaccinated hamsters on days 3, 5, 7 and 14 after challenge with the wild SARS-CoV-2 virus (upper row, **a, b, c, d**). Part (**a)** shows pathohistological changes in the lung were characteristic of injuries in the initial exudative phase of acute respiratory distress syndrome. There is extensive diffuse alveolar damage, atrophy and collapse of the alveoli, desquamation of atypical pneumocytes, peribronchial focal accumulation of pulmonary-associated lymphoid tissue; Part (**b)** shows characteristic atypical cells with pale colored nuclei of different sizes, fibrin exudate and diffusely localized lymphocytes; Part (**c)** shows hemorrhagic necrosis, microthrombi, atypical cells and multinucleated giant syncytial cells, multiple inflammatory cells and fragments of apoptosis. Part (**d)** shows the dominant part of the microstructural elements of the lung parenchyma has already been restored. Bronchiola in a state of recovery with an internal content of red blood cells, microthrombs, hemorrhagic necrosis and diffuse infiltration of lymphoid tissue. The next row has the pathohistological picture of the lung of vaccinated hamsters on days 3, 5, 7 and 14 after control infection with the wild SARS-CoV-2 virus (middle row, **e, f, g, h**). Part (**e)** shows a limited part of the organ parenchyma is subject to surface changes. By the nature of pathohistological changes in the organ, the lungs are in the inflammatory process. In the lumen of only a few alveoli, a small amount of exudate is detected. Part (**f)** shows in the lung, there is an expansion of the zones of diffuse damage to the alveoli. But a significant part of the organ still functions normally, i.e. it has a characteristic microstructure. Part (**j**) shows - the dominant part of the lung parenchyma has a normal microstructure. The connective tissue bases (layers) of the walls of the bronchioles and alveoli have been restored. Part (**h)** shows the hamster’s lungs are externally covered with pleura, formed from the mesothelium and dense elastic connective tissue. Connective tissue partitions divide the lungs into lobes and lobules. The bronchioles of the lungs are covered with a ciliated prismatic or cubic single-layer epithelium. The next row shows the histological picture of the lung of control hamsters on days 3, 5, 7 and 14 (lower row, **i, g, k, l**). Parts (**i, g, k, l)** shows how **t**he characteristic microstructure of the lung is normal. Alveoli, bronchioles, capillaries, respiratory membrane are clearly observed. Stained with hematoxylin and eosin. Increased by x400.

## Discussion

The coronavirus infection COVID-19, caused by the SARS-CoV-2 virus, took on a pandemic character three months after its first appearance and opened a new page for world health in the principles of creating anti-epidemic measures and protecting public health from infectious diseases. It has shown the special need for rapid ways of developing and implementing vaccines against new dangerous infectious diseases. The COVID-19 pandemic has almost stopped the economic life of the planet, caused unique damage to the health of millions of people and claimed hundreds of thousands of human lives (Dong, Du & Gardner, 2020). The current complex epidemic situation and the helplessness of the existing health care system in front of it, forced the need for rapid development of vaccine drugs (Krammer, 2020) and their accelerated introduction into anti-epidemic practice. Due to this situation, a number of countries have not only developed their own vaccines over the past 1-1.5 years, but also successfully conducted three phases of clinical trials (Baden et al., 2021; Polack et al., 2020; Logunov et al., 2021; Voysey et al., 2021).

In Kazakhstan, research on the development of a vaccine against a new coronavirus infection has begun within the framework of a special state scientific and technical program with the appearance of the first cases of the disease in the country in March 2020 on the basis of the SARS-CoV-2 virus isolated from a clinical sample of a sick patient. The research program included the development of five different vaccines: an inactivated whole-virion vaccine based on a virulent virus, a subunit protein vaccine, two recombinant vaccines based on influenza and capripoxvirus vectors, and a live/ replicative vaccine based on an attenuated homologous virus.

In parallel with the ongoing research, at first, the greatest attention was paid to the development of an inactivated vaccine. Because, firstly, the technological basis for the preparation of such a vaccine is more traditional for virological practice, and they have for many decades proven their effectiveness in ensuring biological safety in the fight against such dangerous human diseases as influenza and polio (Katayose et al., 2011; Klein et al., 2020; Krishnan et al., 1983; Grassly, 2014). Secondly, inactivated vaccines, due to the use of inactivated virus, guarantee greater safety. Thirdly, due to the use of a virus containing the entire complex of specific antigens in the composition of whole-virion biomass, the drug provides sufficient immunological effectiveness.

A detailed study of the reproductive properties of the virulent virus, the development of a method for its effective inactivation with the preservation of morphology and antigenic properties, the selection of the correct technological mode of purification of the pathogen allowed us to develop a technology for manufacturing a target vaccine with positive indicators. This paper presents the results of research on the development, evaluation of the safety and effectiveness of inactivated vaccines.

Inactivated vaccines against COVID-19 coronavirus infection are being developed in several countries, including Russia and China. Evaluating the results of a trial in Brazil, China recently reported a low efficacy rate (50%) of its inactivated COVID-19 vaccine (Science, 2021). The decrease in the effectiveness of this vaccine, in contrast to the information previously published about a sufficiently high immunogenicity in China (Gao et al., 2020; Yao et al., 2021), may be associated with the manifestation of a mutated version of the virus (Garcia-Beltran et al., 2021; Wang et al., 2021) or with excessive de-naturation of the virus epitope used in the vaccine during the disturbed inactivation regime, or other factors that may have a negative impact on the immunological effectiveness of the vaccine during production. The negative effect of formaldehyde on the antigenic structure of viral pathogens is confirmed by experimental data of some authors (Metz et al., 2004). For example, when treated with this chemical, the antigenic properties change and the immunogenicity of such vaccines as against viral hepatitis A and B, polio, bovine herpes virus type 1 and influenza decreases when tested on mouse models (Furuya et al., 2010; Tano et al., 2007; Duque et al., 1989; Peterson et al., 1984; Wilton et al., 2014). Taking into account these facts, in order to maximize the preservation of antigenicity, we used the most sparing concentration of the chemical inactivant and the temperature-time mode of inactivation of the virus. As a result, the SARS-CoV-2 virus, inactivated with formaldehyde in the mode selected by us, retained its morphological and structural integrity, which was confirmed by electron microscopic examination, and stimulated the formation of specific antibodies in the body of the laboratory model, which neutralize the virulent virus equally with post-infectious antibodies.

In research and production technologies, beta-propiolactone and gamma rays are also used as inactivants in the inactivation of pathogens of viral diseases (Delrue et al., 2012). At the same time, the researchers note that under the influence of beta-propiolactone, specific proteins of the pathogen undergo significant modification (Delrue et al., 2012). The use of gamma radiation to inactivate viruses is more optimal, since it prevents the formation of free radicals that form toxicity, and reduces the risk of possible changes in the structure of the viral protein. However, there is no detailed information about the use of gamma rays in the preparation of a vaccine against SARS-CoV-2, except in isolated cases (Sir Karakus et al., 2021). In this regard, we did not use this method of inactivation of the virus in our studies.

The safety of the QazCovid-in vaccine candidate has been studied in various laboratory animal species (white mice, guinea pigs, rats), ferrets, and primates (rhesus macaques). As a result, its safety was established on the used animals (the results are not published and are under peer review in Vaccine journal). This article describes the results of studies on the safety, immunological effectiveness and immunity intensity obtained using Syrian hamsters. Thus, according to the literature data, hamsters have unique physiological characteristics, due to which they are well suited for biomedical research as an experimental model (Dutta & Sengupta, 2019) including for viral diseases. It was found that the SARS-CoV-1 virus replicates in these animals (Roberts et al., 2008; Roberts et al., 2005), and, according to the results of recent studies (Sia et al., 2020; Imai et al., 2020; Chan et al., 2020), hamsters, due to their susceptibility to COVID-19 coronavirus infection, are recommended as a biological model for the SARS-CoV-2 virus. Based on these data, we tested the safety assessment of the tested vaccine on Syrian hamsters. When evaluating the safety of the vaccine, there were no visible post-vaccination clinical reactions in all hamsters. In addition to clinical data, indicators of the dynamics of animal body weight, hematological data and biochemical parameters of blood serum were used as indicators of the safety of the tested vaccine. This indicator was evaluated when the experimental animals were administered a 3-fold increased dose of the drug. If the vaccine has any harmful properties for the body, then at such an inflated dose, it should appear pronounced, which can be recorded clinically or by laboratory tests. As shown by the results of studies in hamsters vaccinated with a 3-fold overestimated dose, no deviations in the general condition from the physiological norm were observed. In the animals of the vaccinated group, the live weight gain was higher compared to the animals of the control group (placebo), which indicated the absence of toxicity of the test vaccine.

In preclinical trials to assess the safety of inactivated vaccines against COVID-19, conducted by other researchers, the main focus is on the data of pathological changes detected by pathomorphological and histological methods in parenchymal organs (Gao et all., 2020; Yao et all., 2021; Sir Karakus et all., 2021). At the same time, the data of hematological and biochemical blood tests, which are also informative for assessing the safety of the drug, remain outside the scope of such studies. In this connection, in their studies, they conducted hematological and biochemical blood tests of hamsters in the dynamics of observation. The data of these studies showed that there were no significant deviations from the physiological norm in the studied indicators in both the control and vaccinated groups. The numerical values of the hematological and biochemical parameters of the two groups of animals were also within the physiological norm. All animals during the entire observation period remained healthy and alive without signs of any pathologies.

Thus, the obtained research results fully testified to the safety of the vaccine for hamsters both at the vaccinated human dose, and overestimated threefold.

The main indicators of the immunogenic effectiveness of the vaccine preparation are the factors of humoral immunity in the form of specific antibodies and the resistance of vaccinated animals to the virulent pathogen. Evaluation of the immunity of vaccinated hamsters, conducted by the presence and titer of specific virus-neutralizing antibodies, showed that the QazCovid-in vaccine stimulates the formation of virus-neutralizing antibodies in the body of animals in titers up to 6-8 log_2_, which are detected during the next 6 months (follow-up period) after double immunization with an interval of 21 days. The formation of humoral immunity factors in model animals and humans when using vaccines against COVID-19 coronavirus infection has become publicly available according to existing publications. And these factors are one of the important indicators in assessing the immunogenic reactivity of the vaccinated organism and the immunogenic effectiveness of the vaccine drug used. However, the most reliable way to assess the intensity of immunity formed in response to vaccination with a vaccine is to determine the body’s resistance to disease when infected with a virulent pathogen. Based on this, a number of researchers in their work have shown the protection of model animals immunized with the tested vaccines from coronavirus infection caused by SARS-CoV-2. At the same time, the authors used to evaluate the effectiveness of their vaccine against COVID-19 (Sir Karakus et al., 2021; Roberts et al., 2010; Mohandas et al., 2021; Kandeil, 2021) the level of titer, isolated indicator virulent virus, and histopathological changes developing in the lungs of hamsters were used. In the studies conducted to assess the immunogenic effectiveness of the QazCovid-in vaccine, in addition to the indicators indicated in the publications, we used data on the dynamics of body weight gain, body temperature, clinical indicators and the outcome of the disease in a comparative order in the experimental and control groups of animals. The results of this assessment showed that the vaccinated animals remained at the initial level without weight gain for 7 to 8 days after infection with the virulent virus, and from 9 to 10 days the appearance of weight gain was observed. Among the hamsters of the unvaccinated control group after infection with a virulent virus, in addition to the complete absence of weight gain during the entire observation period, there was a loss of live weight by 20-30 % compared to their initial weight.

The body temperature of the vaccinated hamsters after the control infection with the virulent virus remained within the physiological norm for the entire observation period, and in the control unvaccinated group, 20% of the hamsters had a decrease in body temperature to 34°C. Hypothermia in animals was recorded from 3 to 8 days after inoculation of the virus, and in such cases, the outcome of the pathology was fatal within the same or the next day. Deaths among hamsters used in the experiment for infection with the virulent SARS-CoV-2 virus are not reported in the available publications. According to the authors of such studies (Sia et al., 2020; Imai et al., 2020; Chan et al., 2020), model animals of this species in all cases survived despite the use of high doses of the pathogen for inoculation. In all studies, it was characterized by a decrease in the live weight of animals and the appearance of pathological changes in the lungs of hamsters.

The results of pathomorphological, histological, molecular-genetic and virological studies of lung tissue showed that in all cases, both in the control and experimental groups of animals, the RNA of the virulent virus is detected. When performing virological studies by bioprobe in cell culture, the reproductive virus was detected from the lung tissue of control hamsters, while such a virus was completely absent from the lung tissue samples of vaccinated animals.

During the pathomorphological and histological examination, pronounced pathologies were found in the lungs of the control group of animals from the third day, which persisted until the end of the experiment, and in the lungs of vaccinated animals, such changes of insignificant intensity were noted on 3 to 5 days, which completely disappeared by day 9.

The presence of RNA of the virus and the reproductive pathogen, accompanied by pronounced morphological destruction of lung tissue, indicate the development of acute respiratory disease in control animals as a result of the development of infection after infection with a virulent virus. The absence of a reproductive virus in the presence of RNA of the pathogen and minor morphological pathology in the lungs indicates the reproduction of a virulent virus in the surface epithelial cells of the respiratory system without penetration of the pathogen into the deep layers of tissue, where specific antibodies circulate, preventing the reproduction and spread of the pathogen. The reliability of this pathogenesis is confirmed by the results of tests of inactivated vaccines developed on the basis of related coronaviruses SARS-CoV and MERS-CoV (Bolles et al., 2011; Tseng et al., 2012; Roberts et al., 2010), as well as SARS-CoV-2 (Mohandas et al., 2021; Kandeil, 2021). Also, as in our studies, when infected with a virulent virus of animals vaccinated with a vaccine of these viruses, a mild pathology in the lungs developed.

Investigating the pathogenesis of the disease caused by severe acute respiratory syndrome (SARS) and Middle East respiratory syndrome (MERS) viruses, a number of researchers observed the development of pathology in the lungs associated with antibody-dependent infection enhancement (ADE) (Jaume et al., 2012; Yip et al., 2014; Wang et al., 2016; Yip et al., 2016; Wan, 2020; Perlman & Dandekar, 2005). In addition, there is also information that IgG-class antibodies to SARS-CoV S-protein antigens cause severe macrophage-mediated lung damage in both humans and great apes (Liu et al., 2019). Experiments on rabbits showed that animals re-infected with the MERS-CoV virus by the intranasal method developed pulmonary pathology, accompanied by viremia and severe lung inflammation, despite the presence of specific antibodies formed after the initial infection. In re-infected rabbits, lung damage was more severe than during the primary infection (Liu et al., 2019). In other studies, when animals vaccinated against SARS-CoV (Tseng et al., 2012) or MERS-CoV (Agrawal et al., 2016, p. 2351–2356) were infected with homologous virulent viruses, severe pneumonia developed, despite the high level of specific neutralizing antibodies in the vaccinated animals. Negative consequences from the use of the inactivated virus were also noted in other cases (Bolles et al., 2011, p. 12201–12215).

Based on the above information, we paid close attention to the possibility of such a syndrome when using the QazCovid-in vaccine. However, the results of the studies did not confirm this probability, and the animals vaccinated with the test vaccine, with a control infection with a virulent virus, remained resistant to the disease without developing any visible clinical pathologies in their body. No signs of re-infection or antibody-dependent increased pathology were observed in our studies when animals vaccinated with QazCovid-in were infected with a virulent virus.

## Conclusion

Thus, the candidate vaccine QazCovid-in developed by us has shown itself to be safe and has sufficient protective efficacy against infection with the coronavirus infection COVID-19 caused by the SARS-CoV-2 virus in Syrian hamsters. The immune protection resulting from vaccination with the QazCovid-in vaccine suppressed the replication of the wild virus in the body of vaccinated hamsters, reduced pneumonia and ensured 100% animal survival. Based on the results obtained, the vaccine candidate has been approved for human trials and is currently in the final phase of Phase III clinical trials in volunteers.

## Conflict of interest

The authors state that there is no conflict of interest.

## Author Contributions

Conceptualization, K.Zh., K.Z., L.K. and M.O; planning and conducting experiments, K.S., A.Nr., A.Nk., S.N., O.Ch. B.M., A.I., N.K., S.K., A.K., N.A., Ye.B., M.M., M.A., M.T., data curation, K.Zh, L.K., K.S., A.Nr., O.Ch. Ye.B.; formal analysis, K.Zh, L.K., K.S., A.Nr., O.Ch., M.O., B.Kh., M.K., Ye.A.; funding acquisition, K.Z., Ye.A., and M.K.; investigation, Ye.A., M.K.; methodology, K.Zh., L.K., B.Kh., M.O., K.S., O.Ch.; supervision, K.Z, L.K., M.O., B.Kh; writing-original draft, K.Zh.; writing-review and editing, K.Zh L.K., M.O., K.S., O.Ch., and B.M. All authors have read and agreed to the published version of the manuscript.

## Thanks

The authors of the article express their deep gratitude to the Government, the Ministry of Education and Science, the Ministry of Health of the Republic of Kazakhstan for their assistance in carrying out these research works, as well as to the specialists of the Almaty City Hospital named after I. Zhekenova, personally Deputy Director of the Almaty branch of the RSE “National Center of Biotechnology” Yu. Skiba, Vice-Minister of the Ministry of Health Abishev O. for their assistance in promptly obtaining clinical samples from sick people to isolate the virus, as well as for providing primers for the molecular genetic identification of the SARS-CoV - 2 virus.

The authors of the article also express their gratitude to the staff of the Research Institute of Biological Safety Problems, who provided all possible assistance in these studies.

## Source of funding

The work was carried out within the framework of the scientific and technical program on the topic: “Development of a vaccine against COVID-19 coronavirus infection” (ITN No. 64356/PTF-MES-RK-RL-20) under targeted funding for 2020-2022 with the support of the Science Committee of the Ministry of Education and Science of the Republic of Kazakhstan.

## Footnotes

1 Ventilation mode-15 rpm, CO_2_ concentration 0.04 %, ammonia no more than 0.0001 mg / hour, temperature-19-22 °C, humidity-65-70%.

## Notes

### Competing Interest Statement

The authors have declared no competing interest.

